# A new tool in a toolbox: Addressing challenges in high-throughput microbiota surveys across diverse wild insects

**DOI:** 10.1101/2024.08.26.609764

**Authors:** Mateusz Buczek, Michał R. Kolasa, Monika Prus-Frankowska, Małgorzata Lipowska, Karol H. Nowak, Hamed Azarbad, Diego Castilo Franco, Marzena Marszałek, Tomas Roslin, Anna Michalik, Piotr Łukasik

## Abstract

With their significant effects on the biology of higher organisms, host-associated microbiota has attracted the research community’s attention. The rapid progress in sequencing techniques has greatly facilitated microbial community characterization. However, the most popular surveying technique, marker gene amplicon sequencing, has multiple caveats that are not often addressed satisfactorily, including the uncertainty about the identity of the surveyed wild-caught specimens, variable and sometimes very low abundance of microbes in some samples, or reagent- and cross-contamination. As a result, researchers often obtain incomplete, biased, and sometimes totally incorrect microbial community profiles.

Here, we present a versatile, cost-effective, and high-throughput quantitative multi-target amplicon sequencing workflow for the characterization of host-associated microbial communities, combining laboratory and bioinformatic steps and addressing most of the known methodological issues. Optimized for the study of the microbiota of wild insects, it can be easily adapted for other sample types. Outputs include contamination-controlled data on the absolute abundance and identity of microbes present in insect samples, both at genotype- and OTU-level, as well as host barcodes alongside information on parasite infections. Using 1384 samples from Zackenberg Valley, NE Greenland, we demonstrate the potential of the workflow to study insect and symbiont diversity patterns across a large portion of a diverse natural community.

## 1. Introduction

Microorganisms have played fundamental roles in the evolution of life on our planet, from shaping Earth’s biogeochemical cycles (Falkowski et al., 2008) to affecting many aspects of multicellular organisms biology (Colston & Jackson, 2016; A. E. Douglas, 2016; McFall-Ngai et al., 2013; Rackaityte & Lynch, 2020). Bacteria and unicellular eukaryotes are abundant and diverse in most environments on Earth, including bodies of most higher organisms. Microbial communities associated with diverse eukaryotes have repeatedly been shown to have significant effects on the life histories of their hosts, affecting their populations and short- and long-term evolutionary patterns, with these effects likely reflected within communities (Łukasik & Kolasa, 2024). However, as our understanding of the mechanisms of host-microbe interactions is improving rapidly, the knowledge about microbial community abundance, composition, and significance across and within diverse species that comprise natural communities tends to lag behind (Jansma & El Aidy, 2021; Sieber et al., 2021). This is particularly evident in insects, the most diverse group of animals on Earth. While model systems such as pea aphids, certain *Drosophila* species, and honeybees are well-studied, much less is known about the vast majority of insect clades and species, with many processes and patterns remaining poorly understood (Brown et al., 2023; Chandler et al., 2011; Ferrari et al., 2004; Kwong et al., 2017). Addressing these gaps is critically important and urgent, especially in the context of declining global biodiversity. During times of biodiversity crisis, it is getting increasingly important to target diverse insect communities and understand their symbioses’ diversity, distribution, and spatio-temporal dynamics.

The increasing awareness of the importance of microbiome, combined with technological developments that facilitated their study, resulted in the rapid proliferation of microbiome studies (Thompson et al., 2017). DNA-based methods for microbial community surveys vary dramatically in throughput and unit-cost, as well as information obtained. Currently, microbiome composition is generally studied using two alternative approaches: marker gene amplicon sequencing (metabarcoding) and shotgun metagenomics. Metagenome sequencing can provide higher phylogenetic resolution and a more comprehensive picture of the microbial diversity, host-microbe interactions, and functional relationships between them (Jovel et al., 2016). However, this technology has limitations that restrict its broad application. Microbial strains will only be recorded if their genomic coverage is above a certain abundance threshold - sequencing depth limits the discovery of less abundant but significant microbes. However, the more critical limitations for most users are the relatively high library preparation and sequencing cost and the need for advanced bioinformatic skills and dedicated computational infrastructure for data storage, processing, and analysis. Hence, high-throughput sequencing of marker gene amplicons is the most effective way of broadly surveying microbial community compositions across a large number of samples. Amplicon sequencing is relatively sensitive, fast, and cost-effective, and data analyses are substantially more straightforward than the processing of metagenomic data (Nilsson et al., 2019; Quast et al., 2013; Wang et al., 2007; Yilmaz et al., 2014). Their effective implementation and combination into a cohesive laboratory and bioinformatic workflow, combined with rapidly decreasing sequencing costs, provides an exciting opportunity to address one of the most significant gaps in understanding broad patterns in insect biodiversity and evolution. The method has gained remarkable popularity since its original implementation, arguably peaking with the ambitious microbiota comparison across some 27,000 samples from all types of environments (Meyer et al., 2007; Thompson et al., 2017). However, despite indisputable advantages, amplicon sequencing suffers from several potential limitations and biases that often need to be appropriately addressed. The main limitation refers to the phylogenetic resolution provided by conserved marker genes, which is much less than what can be derived from metagenomic datasets. Additionally, these data provide no functional information despite attempts to create tools for function predictions, such as PiCRUSt2 (G. M. Douglas et al., 2020). In turn, the biases include problems at the stage of DNA extraction, library preparation, and sequencing, including reagent- and cross-contamination, sequencing errors, and chimera formation (Knight et al., 2018).

The selection of appropriate bioinformatic tools for data processing, including accurate taxonomic identification of microbes, data filtering and binning, and biologically relevant interpretation of results, can be another challenge, particularly for non-model biological systems. However, one of the method’s main limitations is the amplification bias against certain taxonomic clades caused by different primers. None of the broad-spectrum primers targeting different rRNA variable regions is truly universal, with differences in amplification efficiency among microbial clades that make up the community, with some promoted and others underrepresented, or even excluded entirely, by certain primer sets (Bukin et al., 2019; Klindworth et al., 2013; Walters et al., 2015; Willis et al., 2019). This led to the introduction of modified, more degenerate primer sets, like those adopted by the Earth Microbiome Project (EMP), probably the largest amplicon-based study to date. Still, the V4 primers currently recommended by the EMP perfectly match only 88.2% of bacterial sequences within the Ribosomal Database Project on that date, with some taxa covered much less well than others. While imperfect primer matches do not preclude amplification, primer biases remain a concern in microbiome surveys. Therefore, an essential prerequisite for broad microbiome studies is developing a method capturing the large majority of microbes present in insect samples, including strains divergent from those in reference databases and barcode hosts and parasites that may be another source of microbes (Kolasa et al., 2023).

To address these challenges, we developed, optimized, and successfully implemented a protocol for quantitative multi-target amplicon sequencing - a versatile tool combining amplification of marker genes: host COI and bacterial 16S rRNA, which can be easily extended to other targets, e.g., fungal ITS sequences. We show that amplicon sequencing can be a powerful and effective way of capturing diverse insect-associated bacteria and demonstrate its utility in a diverse wild-caught community from Greenland.

## 2. Methods

### 2.1 Validating laboratory solutions

#### Overall strategy

Further in the Methods (section 2.3), we outline the final version of the laboratory protocols for quantitative multi-target amplicon sequencing workflow, which are presented in detail on protocols.io: <u>dx.doi.org/10.17504/protocols.io.36wgq351ylk5/v1</u>. Here, we describe how we have tested different steps of the protocol, arriving at specific solutions. Generally, we used replicate specimens of different insect species originating from laboratory cultures or relatively homogenous wild populations (Suppl. Table 1), and used different methodological solutions for one of the laboratory steps while keeping the other constant. We summarize the details in further sub-sections, with details of each assay in Supplementary Material.

##### 2.1.1 Testing alternative DNA extraction approaches

Initially, we assessed the effects of alternative DNA extraction methods on insect microbiome profiles by comparing microbial community profiles among samples processed using commercial kits (Qiagen DNeasy Blood and Tissue, Qiagen DNeasy Powersoil) and a custom procedure described in the final protocol. Using those three methods, we extracted DNA from 6 different groups of insects (Orthoptera: *Gryllodes sigillatus*, Diptera: *Drosophila teisseiri*, Hymenoptera: *Cephalotes varians*, Blattodea: *Ectobius lapponicus*, Hemiptera: *Macrosteles laevis*, Coleoptera: *Otiorhynchus equestris*) and microbial standard (75ul ZYMO Microbial Standard D6300). Using two-step PCR, we prepared amplicon libraries for three target genes (COI and bacterial 16S V1-V2 and V4 regions) using Qiagen Multiplex polymerase (see Supplementary Materials).

For planthoppers used for bacterial primers validation, we used two different DNA extraction kits: for putative fungus-associated species - Genomic Mini AX Yeast Spin kit (A&A Biotechnology) and Sherlock AX isolation kit (A&A Biotechnology) for the remaining species.

##### 2.1.2 Bacterial primers validation

We tested to what extent three alternative primer pairs targeting different regions of bacterial 16S rRNA gene (V1-V2, V3-V4, and V4) affect our ability to detect symbionts in 18 planthopper species (Hemiptera: Fulgoromorpha), a clade known to host highly specialized, rapidly evolving bacterial or fungal heritable symbionts (Michalik et al., 2023), likely constituting a particularly challenging target for “universal” bacterial primers. Specifically, we used metagenome-recovered ribosomal RNA as a reference against which amplicon datasets for multiple target regions, obtained for the same DNA samples, were compared and calibrated. Amplicon libraries were prepared using primers specific to V1-V2, V3-V4, and V4 regions of the 16S rRNA gene (Suppl. Table 2.) and sequenced on Illumina MiSeq using v3 2x300bp kit at the Institute of Environmental Sciences of Jagiellonian University (Kraków, Poland). Metagenomic libraries were prepared using the NEBNext Ultra II DNA kit, pooled and sequenced on 2x150 bp HiSeq X lane by NGX Bio (San Francisco, CA, U.S.A.).

##### 2.1.3 Library preparation approaches comparison

We have tested alternative indexing schemes for amplicon-library preparation: Adapterama, Andersson, Meyer & Kircher (Glenn et al., 2019; Marquina et al., 2019; Meyer & Kircher, 2010). Using simple Taq polymerase, we amplified two target genes separately using primers described in previous sections: COI and V4 region of 16Sr DNA. Each PCR product was cleaned using magnetic beads and then diluted 1000x or 1000000x times. Next, we performed second-step PCR (indexing) using three pairs of primer (Adapterama, Andersson, Meyer & Kircher), three template concentrations (diluted: 1x, 1000x, 1mlnx), and two High Fidelity polymerases (Q5 and KAPA HiFi).

##### 2.1.4 Multiplex PCR optimization and High-Fidelity polymerases validation

We have tested the ability of different polymerases to amplify target genes. Additionally, we checked how some of them performed in multiplex PCR. We run PCR for four target genes using nine different polymerases for seven distinct insect DNA samples. Next, for an additional six insects and microbiome standard, we run PCR for three target genes separately and in multiplex, using HiFi polymerases (KAPA and Q5) and Qiagen multiplex.

We tested the performance of High Fidelity polymerases and Qiagen Multiplex in second PCR (indexing). Since only Qiagen multiplex performed well during multiplex PCR, we used the PCR product of this reaction to check if using high-fidelity polymerases (KAPA and Q5) in the second PCR affects the quality of obtained sequences. Furthermore, using Qiagen Multiplex polymerase, we performed three separate PCRs for three target genes (COI, 16S_V4 16S_V1-V2) separately. Next, we mix three PCR products: COI, 16S_V4, 16S_V1-V2 in proportion 1:1:1 and run a second PCR using three polymerases: Qiagen, KAPA, Q5.

Additionally, we tested whether the type of polymerase affects the recovery of all insect sequences from the mock community sample. We isolated DNA samples using our custom method from a mock community sample containing 23 insect species. Using isolated DNA, COI, and 16S rRNA V4 region primers and Qiagen Multiplex, we performed PCR in three replicates and two cycle regimes (15 or 25 cycles). Next, using four polymerases (Phusion, KAPA, Q5, and Qiagen Multiplex), we performed a second PCR (indexing) using 15 or 5 cycles, respectively.

##### 2.1.5 Quantification approach validation

To calculate the absolute abundance of the insect microbial community, we used an artificial construct (spike-in) targeting the V4 region of the 16S rRNA bacterial marker gene (Tourlousse et al., 2017). The primer binding sites matched the primers used for bacterial metabarcoding (Apprill et al., 2015), and the remainder of the sequence was replaced by a random DNA sequence that does not resemble any known sequence deposited in GenBank while maintaining GC content similar to that of bacterial DNA. Spike-ins were synthesized and inserted into the pEX-a258 plasmid vector (Eurofins) and then transformed into competent *Escherichia coli* TOP10F cells (A&A Biotechnology). The *E. coli* cells with integrated plasmid were then selectively grown overnight on LB-ampicillin medium (50 mg/ml), and finally, plasmids were extracted using a Plasmid Miniprep DNA purification kit (Eurx). Post-extraction, the plasmids were linearised and quantified. Subsequently, appropriate spike-in dilutions were prepared based on the DNA aconcentration. The primary objective was to add an equal number of spike-in copies to each homogenate sample immediately before DNA extraction (Suppl. Table 1).

We used 290 specimens belonging to 24 species (Suppl. Table 1) to extract DNA with added spike-ins after the homogenization. Next, DNA was used to prepare amplicon libraries targeting COI and 16SV4 marker regions, and a subset of one specimen per species was used to prepare metagenomic libraries. Amplicon libraries were processed using a set of customized scripts (see section 2.2). Whereas metagenomic libraries were analyzed using established tools. First, adapters were trimmed using Trimgalore (Krueger, 2012) and then contigs were assembled using Megahit (D. Li et al., 2016). Next, we used bowtie2 to create a reference and map raw reads against it (Langmead & Salzberg, 2012), followed by converting .sam files into .bam files using Samtools (H. Li et al., 2009). Finally, we used Anvio (Eren et al., 2015) to extract 16S sequences (including quantification spike-ins) from metagenomic libraries and calculated absolute abundance similarly to in amplicon libraries.

### 2.2 Validating bioinformatic approaches

#### 2.2.1 Overall strategy

Multitarget amplicon data was analyzed using a set of custom Python scripts combining already established tools for microbiome analysis. A detailed description of each step is provided in the GitHub repository: https://github.com/Symbiosis-JU/Proof_of_Concept. Here, we briefly describe each step of the analysis.

#### 2.2.2 Splitting sequences into bins

As mentioned in the previous section, each library consists of a mixture of target sequences such as host COI, bacterial 16S V4 and V1-V2. The splitting script recognizes the sequences of primers used for each of the target regions (separately for FASTQ R1 and R2 files) and splits them into corresponding bins. Simultaneously, as we use informative indexes, the script removes potential cross-contamination. Each position in the PCR plate is assigned to a specific indexing scheme. Hence, if the library consists of any sequences that are characteristic of the position of different samples, they’ll be removed. After running the script, the user is provided with directories corresponding to each target region with cross-contamination-free R1 and R2 FASTQ files.

#### 2.2.3 Joining pair-end reads, quality control, denoising, zOTU/OTU tables outputting

This part is the core of the analysis. First, the script uses PEAR (Zhang et al., 2014) to join R1, and R2 reads based on the specifications provided by the user: the minimum and maximum length of amplicon after joining, the minimum overlap between R1 and R2 sequences and the minimum Phred (quality) score. Next, VSEARCH (Rognes et al., 2016) converts FASTQ to FASTA files, dereplicates them, and denoises each library separately. This step is extremely important, as denoising all libraries together can lead to the loss of biologically relevant biodiversity (Kolasa et al., 2023; Prodan et al., 2020) It then uses USEARCH (Edgar, 2016) for OTU-picking and chimera removal. Finally, it assigns taxonomy classification for each of the sequences based on customized databases such as MIDORI (version GB 239) (Leray et al., 2022), SILVA (version 138 SSU) (Quast et al., 2013) and outputting zOTU and OTU tables.

#### 2.2.4 Decontamination and microbiome absolute abundance calculating

The next script uses the output from the previous step, as well as information about various types of negative controls (extraction, PCR and indexing negative controls) and the type of artificial spike-in added to decontaminate and quantify bacterial data. It produces four tables: Table_with_classes, which is an input table with each bacterial zOTU assigned to one of the categories: extraction spike-in, PCR spike-in, non-bacteria, extraction contaminant, PCR contaminant, symbiont, other (zOTU classified as symbiont, but with the maximum relative abundance in any of the sample libraries <0.001% [the threshold indicated by the user]); statistic tables with statistic for each library containing the read and abundance information about zOTUs classified as symbionts, contaminants and spike-ins as well as the ratio between symbionts’ reads and spike-in’s reads; and finally decontaminated zOTU and OTU tables with information about each zOTU’s taxonomic classification and OTU affiliation, marker gene sequence, and total number of reads across libraries. First, it uses taxonomic classification to delete all zOTUs assigned as chloroplast, mitochondria, Archaea, Eukaryota or chimera. Next, based on the negative controls list, the maximum abundances of each of the zOTU in each of the negative controls and experimental samples are calculated. Based on the thresholds provided by the user, each zOTU is assigned to the category of symbiont or contaminant. Finally, it calculates the proportion between artificial spike-in reads and sequences assigned as symbionts. This ratio, combined with the information about the proportion of insect homogenate taken for DNA extraction and the number of spike-in copies added at the extraction step, allows us to calculate the absolute abundance of bacteria for each library.

#### 2.2.5 Barcode generation and COI data filtering

The next script, ICRC (Insect COI Reads Classifier), uses COI zOTU/OTU table generated in step 2.2.3. It has two functions: finding the barcode sequence for each library and classifying reads into multiple categories. Two files, FASTA and classification table, are generated accordingly. The FASTA file contains barcode sequences (most abundant, non-bacterial zOTU). The classification table contains a table of reads counts divided into multiple categories, such as barcode, secondary barcode, taxonomy groups (e.g., *Wolbachia*, plants), potential parasitoids, and cross-contamination. A detailed description is provided in the supplementary materials.

### 2.3 Final protocol

The proposed protocol for high-throughput microbiota characterization involves essential laboratory steps, including DNA extraction, Illumina amplicon library preparation, sequencing, and subsequent bioinformatic analyses enabling the determination of insect host identity, reconstruction of bacterial diversity, and bacterial quantification. The key steps of the protocol are illustrated in Fig. 2.

**Fig. 1.**
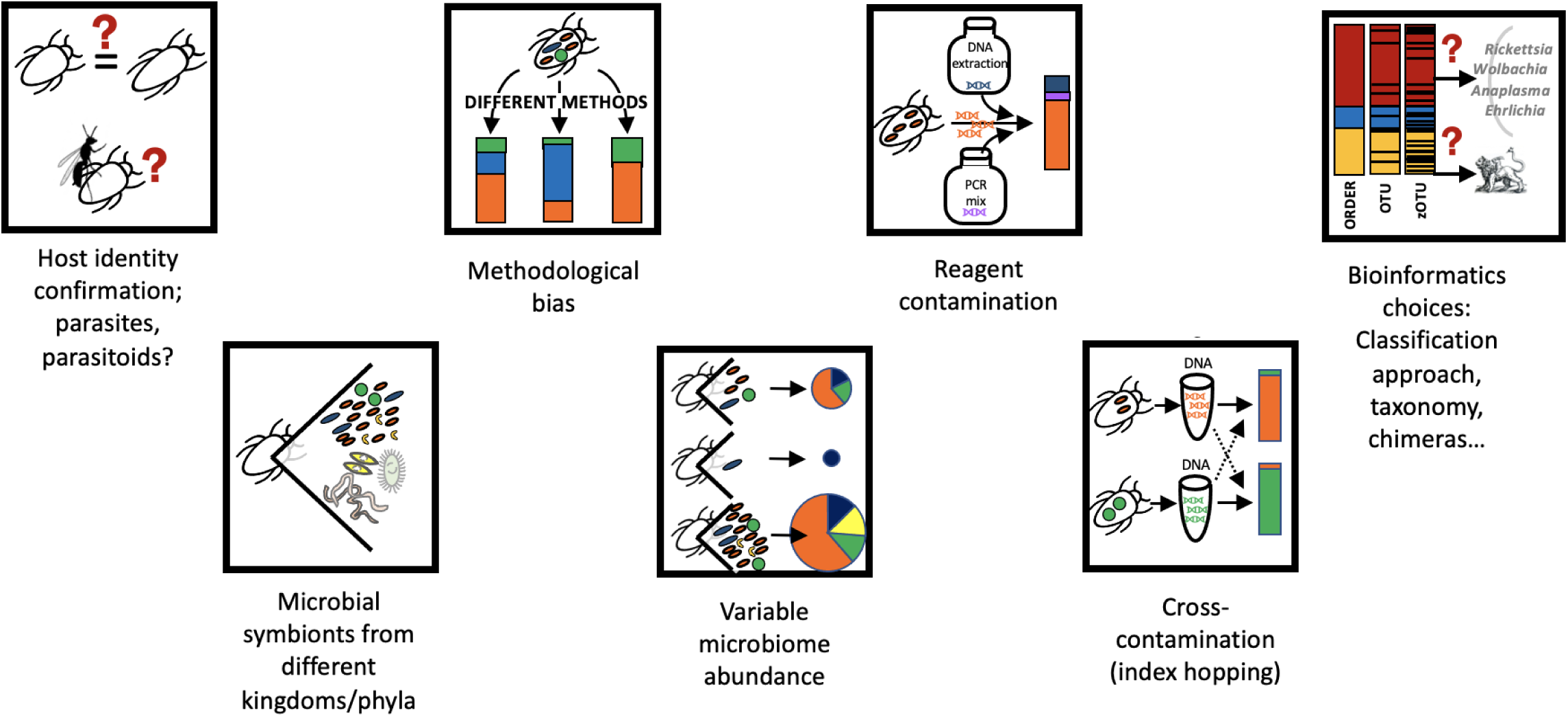
Some of the conceptual and methodological challenges in amplicon-based microbiome studies across diverse wild hosts.

**Fig. 2.**
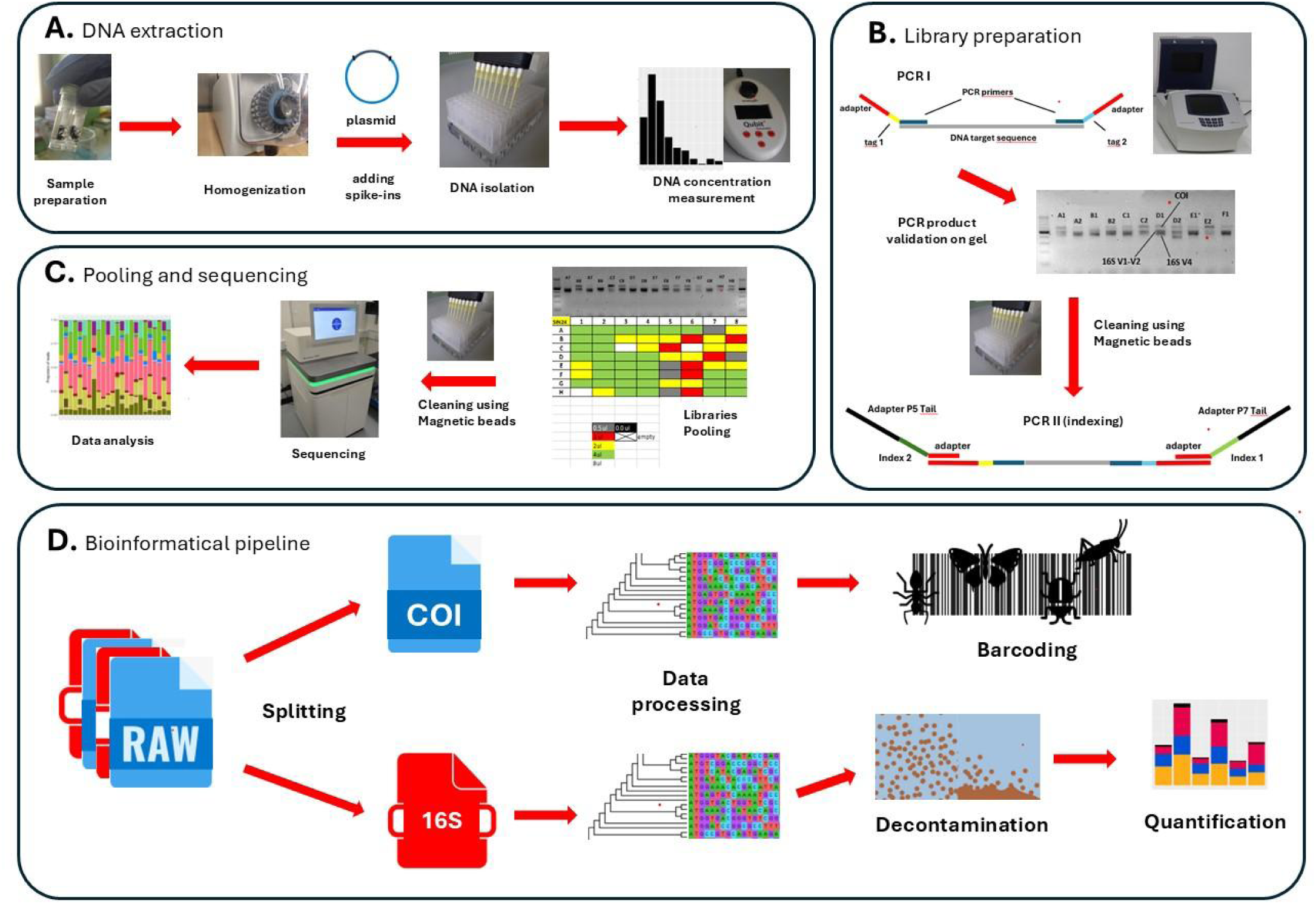
The schematic overview of the proposed method/protocol workflow.

### 2.4 Validating protocol on wild-caught insects

To test the utility of the workflow and assess labour intensity and cost, we used a collection of insects obtained during a sampling trip to Zackenberg Valley in NE Greenland in 2021, where the relatively limited set of ca. 1300 insect species makes it plausible to sample and reconstruct community-level host-microbe patterns comprehensively. With its low insect diversity and very comprehensively described ecological interactions, Greenland has a strong potential to be used as a model of how natural communities have been shaped and how they respond to climate challenges.

Insects were collected using Malaise traps and entomological nets. They were stored in 96% ethanol and stored at −20℃ till processing in Poland. Samples were screened from each location and collection date, and individual insects were picked to represent the majority of morphological diversity in each sample. 1583 insects were chosen for processing (Suppl. Table 3). Each individual was assigned to a size category, homogenized and DNA was extracted according to our custom protocol. Next, amplicon libraries for COI and 16S rRNA V4 regions were prepared and sequenced on Illumina NovaSeq 6000 at the National Genomics Infrastructure (NGI) in Stockholm, and raw reads were processed with our customized bioinformatic pipeline (see previous sections of Methods).

## 3. Results

### 3.1 Testing protocols

#### 3.1.1 Testing alternative DNA extraction approaches

The three tested DNA extraction methods showed no significant differences based on PCR band intensity. All target genes were successfully amplified, with consistent patterns within sample triplets (Suppl. Fig. 1). Marker gene reads composition was also consistent within the sample triplets (Suppl. Fig. 2 and Suppl. Fig. 3). However, bioinformatic analyses revealed that the DNeasy Blood and Tissue kit (Qiagen), failed to extract spike-ins successfully as no spike-in reads were obtained for samples processed using this kit. In turn, the Genomic Mini AX Yeast Spin kit (A&A Biotechnology) introduced a significant contaminant *Cellulosimicrobium* - a microbe used for the commercial production of the lyticase enzyme, as reads corresponding to this taxa dominated all the samples processed with this kit (Suppl Fig 4).

#### 3.1.2 Bacterial primers validation

For each of the 18 experimental planthopper species, we obtained amplicon data for three targeted 16S rRNA regions. After removing singletons and chimeras, we obtained an average of 25,688 reads for the V1-V2 region, 10,986 reads for the V3-V4 region, and 31,462 reads for the V4 region. Post-denoising and decontamination, we identified 229 zOTUs for V1-V2, 222 for V4, and 102 for V3-V4 across all samples. In turn, using PhyloFlash, we recovered from all metagenomes 77-full-length bacterial 16S rRNA sequences. Among them, we identified bacteria representing genera known as insect symbionts (e.g., *Sulcia*, *Vidania*, *Sodalis*) and bacteria that appear to have originated from laboratory reagents (e.g., *Cellulosimicrobium*) (Michalik et al., 2023; Suppl. Fig. 4). We compared the amplicon-derived zOTUs against 16S rRNA sequences from metagenomes for the same DNA samples. This comparison enabled us to confirm whether amplicon sequencing captured all genotypes identified in the metagenomes and identify additional microbes found in the amplicon data but not in the metagenomes, along with their likely origins.

We found that the large majority of reads in the planthopper libraries represented zOTUs that matched full-length metagenomic sequences for the same planthoppers. Conversely, in most cases, the most abundant amplicon zOTUs matched the metagenomics sequences precisely (Suppl. Fig. 4) and Supplementary Table 4). In total, out of 77 16S rRNA sequences assembled from 18 metagenomes (including real insect associates and apparent contaminants), V1-V2, V3-V4, and V4 regions have retrieved 69, 72, and 68 sequences, respectively (Suppl. Table 4). However, considering only those 16S sequences regarded as the real insect associates and excluding contaminants, in almost all cases, the V3-V4 and V4 fragments detected 100% of all the microbes determined with metagenome-based sequencing. There is only one exception - in the V4 dataset for *M. pruinosa*, we failed to observe *Wolbachia* symbiont. In turn, the V1-V2 region failed to detect *Vidania* in the case of *Asiraca clavicornis* and *Stenocranus major* (Suppl. Fig. 4). The relative abundance estimates for different symbiont taxa were also highly correlated; however, there were notable exceptions (e.g., estimated *Vidania* abundance in *C. pilatoi* based on V4 region compared to data from other regions and the metagenome).

Microbial zOTUs other than the “verified” symbionts also present in metagenomes made up 2,8% of the dataset on average. Among them, putative DNA extraction contaminants (e.g., *Cellulosimicrobium* derived from the kit dedicated to fungal DNA extraction) dominated. Additionally, in amplicon datasets for all libraries, we observed bacteria that were absent in metagenomes, including some known PCR contaminants (e.g., *Brachybacterium*) (for details, see section 3.1.6).

Another challenge with amplicon sequencing data analysis is selecting an appropriate database for the taxonomic classification of zOTUs sequences. In our analyzed dataset, taxonomic classification and binning of zOTUs using a standard approach against the SILVA v.138 database only partially overlapped with results for full-length sequences. Notably, some sequences that matched full-length metagenomic sequences perfectly were left unclassified, and in other cases, zOTUs that corresponded to different regions of the same full-length sequence were classified differently. This issue was particularly evident with *Vidania*, an extremely rapidly evolving betaproteobacterium that was repeatedly misidentified as betaproteobacterial endosymbionts of related hemipterans, such as *Nasuia* or *Zinderia*.

#### 3.1.3 Library preparation approaches comparison

The results presented in Suppl. Fig. 5 show that the Meyer & Kircher indexing scheme is prone to form primer dimers (at least in the case of low-concentration templates). On the other hand, Andersson and Adapterama managed to amplify all the samples. However, Adapterama gives slightly stronger bands than Andersson, and the difference is not caused by the low primer concentration (see primer leftovers marked on the figure by yellow circles in Suppl. Fig. 5).

#### 3.1.4 Multiplex PCR optimization and High-Fidelity polymerases validation

The only polymerase that amplified all target genes in all the samples was Qiagen Multiplex (Fig. 3 and Suppl. Fig. 6). Lack of bacterial 16S signal in some of the samples amplified by Qiagen Multiplex results from the low microbial signal in those samples (reflecting the biology of those species). The low performance of most of the polymerases in amplifying COI targets is probably caused by higher primer degeneration. Only Qiagen Multiplex deals with multiplex PCR (Suppl. Fig. 6). Hence, with a two-step PCR approach, Qiagen Multiplex seems to be the only viable option. Moreover, no other polymerase was able to consistently amplify different marker genes separately.

**Fig. 3.**
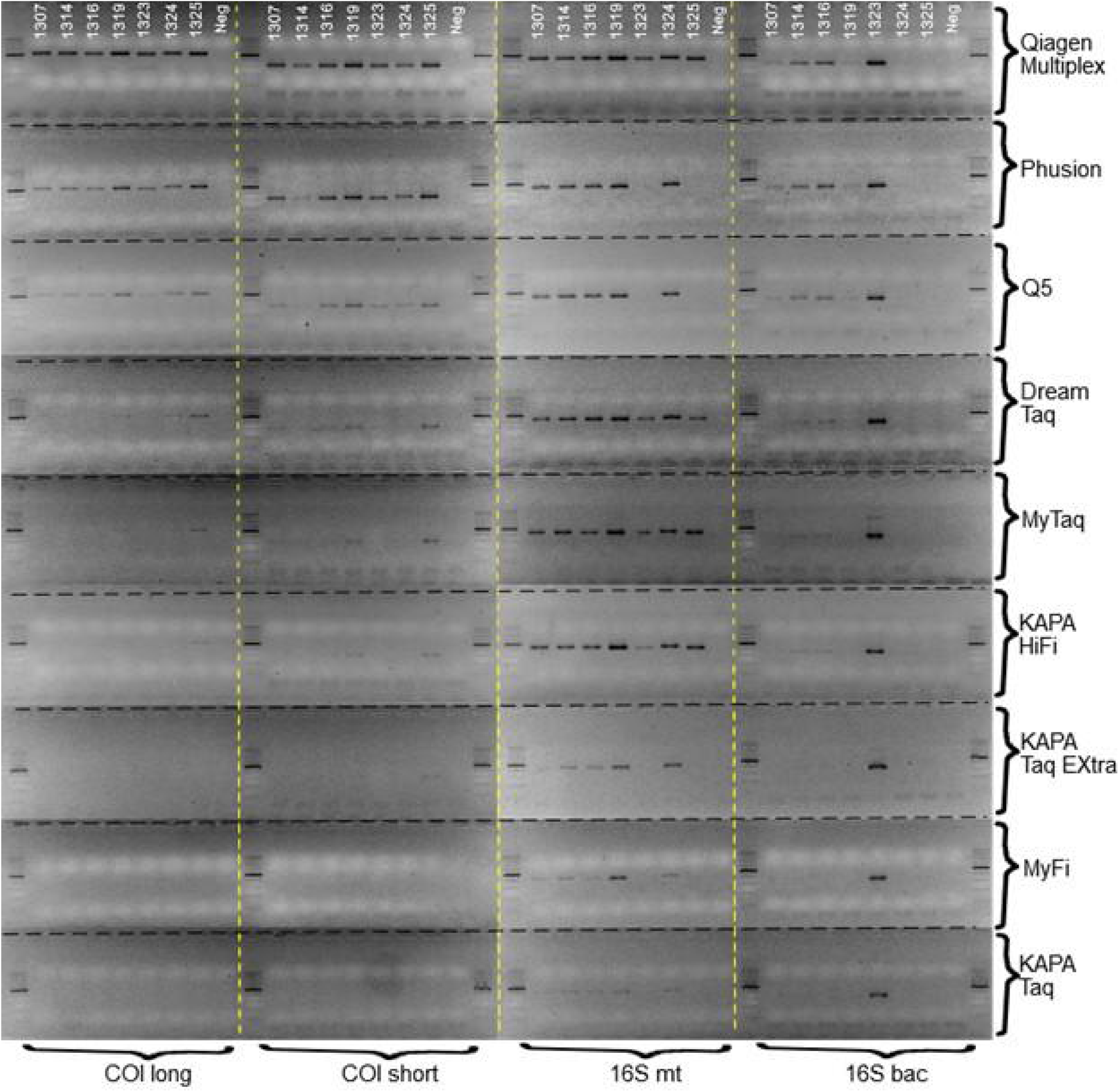
Comparison of different polymerases to amplify various marker genes (COI long fragments, COI short fragments, 16S mitochondrial and bacterial 16S V4).

After the second PCR (indexing), marker genes reads composition was consistent for both HiFi polymerases and Qiagen Multiplex (Suppl. Fig. 7). As for an error rate (counted as a proportion of the most abundant OTU sequence), the differences between polymerases are negligible (Suppl Table 5). Additionally, as visualized at Suppl. Fig. 8, polymerase Q5 preferentially amplifies shorter fragments, as the proportion of COI product is noticeably smaller compared to KAPA and Qiagen.

Only KAPA and Qiagen Multiplex manage to recover signals of all mock community insects and perform much better than Phusion and Q5 (Suppl. Fig. 9). Additionally, it looks like we get slightly higher insect species recovery using scheme 15 cycles in PCR I and 15 cycles in PCR II (indexing). Furthermore, Qiagen Multiplex is much cheaper than any of the three tested HiFi polymerases.

#### 3.1.5 Decontamination and Quantification

Our pre-decontamination zOTU table comprised 315 libraries, including sample and control libraries. The total number of reads was 6,358,372 and ranged from 18 (in one of the blank samples) to 105,156, while excluding blank samples, they ranged from 653 (for one of *Dacnusa sibirica* samples) to 105,156 (for *Shelfordella lateralis*) with a mean value equal to 21,037 reads (Suppl. Table 6). After decontaminating using our custom script and excluding blank samples, the number of reads was equal to 5,042,343 and ranged from 0 (for one of

*Dacnusa sibirica* samples) to 93,341 (for one of the *Shelfordella lateralis*) with mean value equal to 17,387 reads (Suppl. Table 7). In total, 109145 reads were assigned as non-bacteria or Extraction/PCR contaminants across sample libraries, which constituted 1.79% of total reads. The most abundant PCR contaminant zOTU, was taxonomically assigned as *Brachybacterium* (38,205 reads, 0.63% relative abundance across all libraries), while the most abundant extraction contaminant was assigned as *Klebsiella* (4918 reads, 0.08% relative abundance across all sample libraries). Non-bacteria (zOTUs assigned as chimera, mitochondria, Eukaryota, chloroplast, and Archea) accounted for 0.86% of overall relative abundance with 52,450 reads across all libraries (Suppl. Table 6).

After decontamination, we proceeded to calculate the absolute abundance of microbial community in amplicon libraries based on the symbiont: extraction spike-in ratio (calculated with our custom Python script), the number of extraction spike-in copies added to a homogenate, and the fraction of homogenate taken for DNA extraction. For 62 out of 290 samples (processed with DNeasy Blood and Tissue kit), we were unable to calculate absolute abundance due to a lack of extraction spike-in reads in the libraries (Suppl. Table 8, Fig. 4). Across all studied species, the highest absolute abundance characterized *Apis mellifera* (min log. 8.67, max log. 9.57, mean log. 9.13), whereas the lowest was calculated for *Dacnusa sybirica* (min log 3.19, max log. 4.94, mean log. 4.44) (Suppl. Table 8, Fig. 4).

**Fig. 4.**
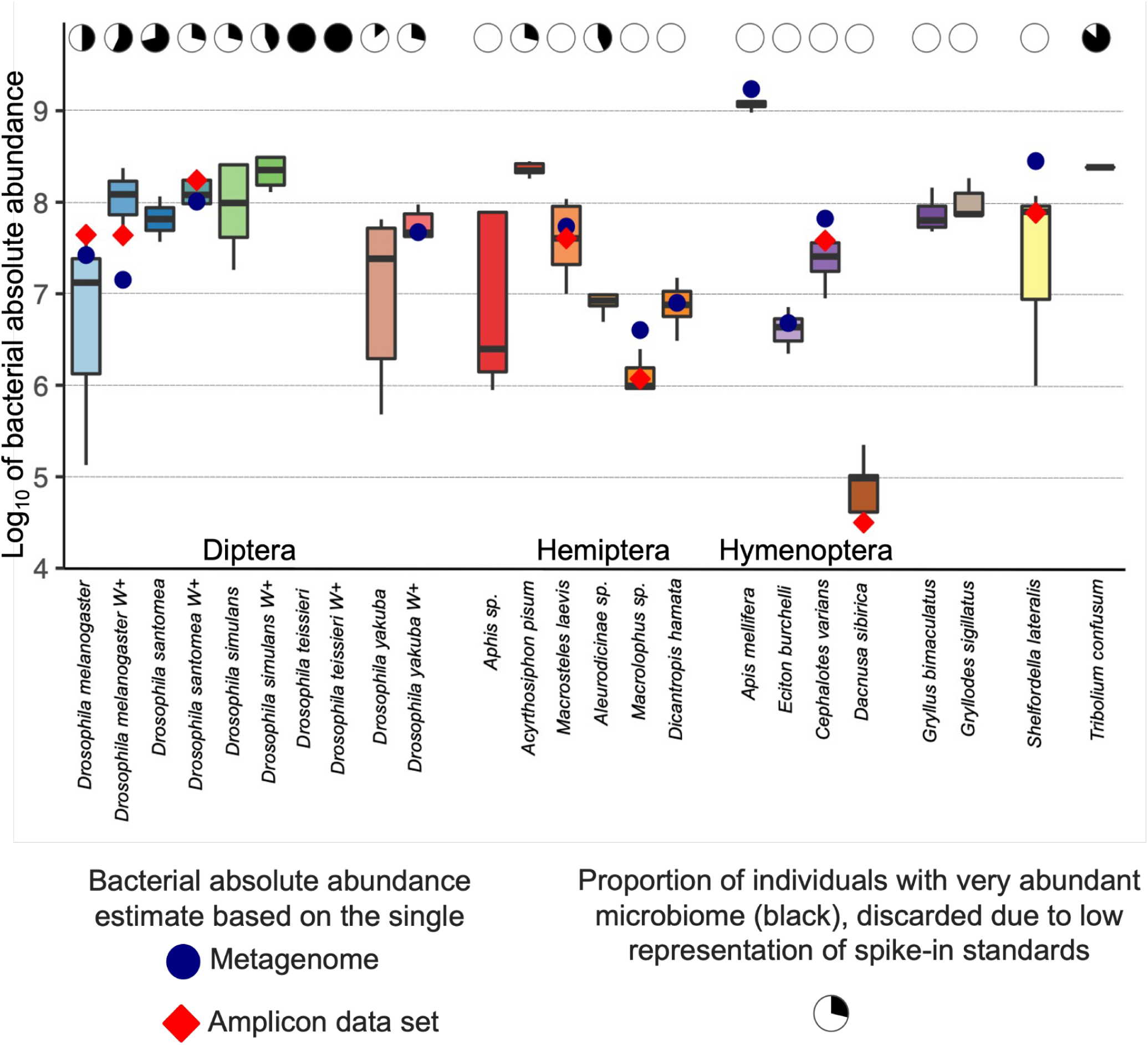
Series of box plots representing amplicon-based absolute abundances for the specimens from the different species grouped by order. The blue dot and red diamond represent the decimal logarithm of absolute abundance (metagenomic and amplicon-based, respectively) for the same specimen. Specimens we could not confidently calculate absolute abundance (no or too few extraction spike-in reads in the library) were excluded from the analysis.

Next, we calculated absolute abundance for a subset of 24 metagenomic libraries. For 12 libraries, we were unable to reconstruct the bacterial 16S gene or quantification spike-in reads (Suppl. Table 8.). For the rest of the samples, quantification based on amplicons and metagenomes was congruent (Suppl. Table 8, Fig. 4). The highest difference between amplicon and metagenomic quantification was observed in *Aphis sp.* (amplicon log. 7.9, metagenomic log. 6.68), the lowest in *Dicanotropis hamata* (amplicon log. 6.89, metagenomic log. 6.9) (Suppl. Table 8).

### 3.2 Validation with natural insect community

First, all 1650 libraries were separated into bins corresponding to marker regions and removal of potential cross-contamination.

#### 3.2.1 Cytochrome oxidase subunit I data

After joining forward and reverse reads, keeping only high-quality reads (Phred score >30), and singleton removal, we obtained high-quality COI data for 1572 samples, including negative and positive samples. Overall, we obtained 54,211,877 COI reads for all libraries (min. 2, max 119,497, mean 34,486) (Suppl. Table 9). Next, we proceeded with our script, which was responsible for barcode recovery. Applying a minimum number of reads for barcode equal to 1000 and removing positive and negative samples, we ended up with barcodes for 1384 samples (Suppl. Table 10). The reads representing barcodes ranged from 1008 to 117,684 (mean 36,546). The vast majority (1084) of barcode zOTUs represented >0.99% of abundance within the library. The most abundant insect family was Chironomidae, with 790 specimens, followed by Culicidae, with 138 specimens. In 194 COI libraries, we found the zOTUs classified as bacteria (mainly *Wolbachia*) (Suppl. Table 10, Fig. 5).

**Fig. 5.**
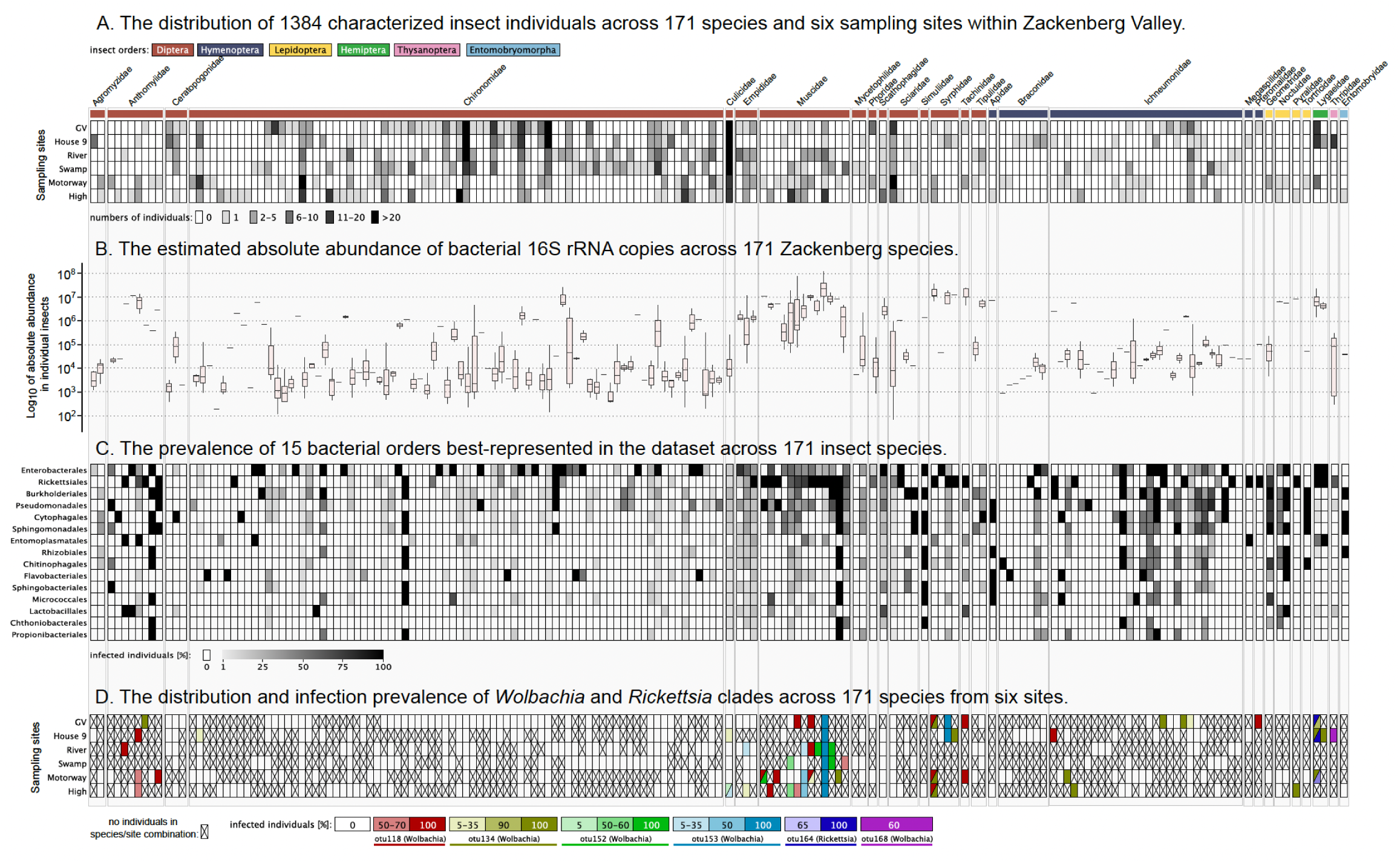
Distribution of insect species and their microbiomes at six locations in the Zackenberg Valley in North East Greenland. Host barcode data indicate that we have captured about 143 species, or over half of the insect species previously recorded in Zackenberg Valley, in addition to 28 species new to the area. These insect species differed in their prevalence across the sampled habitats (A), as well as in terms of their microbiome. The total abundances of bacteria hosted by individual insects varied among samples by more than five orders of magnitude (i.e., 100,000-fold), but were low in most individuals (B). The prevalence of bacterial clades differed among insect species (C). At finer taxonomic scales, some bacterial clades, including facultative endosymbionts *Wolbachia* and *Rickettsia*, were detected in multiple insect species from different families (D), and their prevalence within some species differs across habitats.

#### 3.2.1 Bacterial 16S rRNA V4 data

The raw pre-decontamination dataset consisted of 1655 libraries (including various negative and positive samples) with a total of 40,929,737 reads (min. 2; max 159,074; mean 24,731) (Suppl. Table 11). After the decontamination and removal of positive (37) and negative (90) samples, we decided to remove an additional 144 samples absent in the COI database and ended up with 1384 samples with high-quality COI and 16S rRNA datasets (Suppl. Table 12). The total number of bacterial reads across the final dataset reached 33,829,935. In the final dataset, zOTUs assigned as PCR contaminants had a total of 1,379,360 (4.08% of total abundance). The most abundant taxa in this category were *Brachybacterium* (779,541 reads, 2,3% of relative abundance), followed by *Polaromonas* (421,344 reads, 1.24% of relative abundance). The rest of the less abundant 191 zOTUs assigned as PCR contaminants in total made 0.54% of relative abundance with 178,475 reads (Suppl. Table 12). Extraction contaminants, on the other hand, were significantly less abundant. In total, 156 zOTUs classified in this category made 1.96% of total abundance with 66,235 reads. The two most abundant zOTUs were classified as *Spingomonas* (13,377 reads; >0.04% of abundance) and *Corynebacterium* (7859 reads, 0.02% of abundance). 7514 zOTUs were classified into the non-bacteria category with 1,088,804 reads and 3.22% of relative abundance. The three most abundant zOTUs were classified as chloroplast with a summed relative abundance of 0.68% (230,658 reads). The fourth and fifth most abundant non-bacteria were classified as mitochondria with a summed relative abundance of 0.37% and 154,500 reads (Suppl. Table 12). In the final dataset, we obtained 37,641 zOTUs grouped into 9497 97% similarity OTUs (Suppl. Table 13). The ten most abundant OTUs classified as Morganellaceae (1), *Wolbachia* (2), *Rickettsia* (1), *Schneideria* (1), *Spiroplasma* (2), *Rhanella* (1), *Pseudomonas* (1), and *Cardinium* (1) made up 77.88% of all reads across all libraries (min. 171,496; max 2,594,817; mean 949,928) (Suppl. Table 13). The most abundant OTU in read number was taxonomy classified to Morganellaceae family characteristic for chironomids (2,594,817 reads, 21,27% of all reads). This is unsurprising since chironomids were the most abundant taxonomic group in the dataset. This OTU consisted of 50 zOTUs, with 13 exceeding 1000 reads across all libraries (min. 2, max. 1,610,671, mean 51,896) (Suppl Table 13). The second most abundant OTU was classified as *Wolbachia* (2,558,811 reads, 20.98% of all reads). It consisted of 116 zOTUs, with 14 exceeding 1000 reads across all libraries (min. 2, max 728,765, mean 22,059). In total, we obtained 23 OTUs classified as *Wolbachia* with variable read numbers (min. 2, max 255,881; mean 153,124; total 3,521,856), and 6 of them exceeded 1000 reads across all libraries. In total, *Wolbachia* reads made up 28.87% of the dataset.

The lowest absolute abundance was observed among chironomids (max. log. 7.43, min. log. 1.64, mean log. 4.01). In contrast, the highest was found in the Muscidae family (max. log. 8.1, min. log. 4.67, mean log. 6.69) (Suppl. Table 14, Fig. 5).

## 4. Discussion

Over the last two decades, our understanding of microbiome diversity, composition, and its role in various biological systems has significantly advanced (Lloyd-Price et al., 2017). With its increasingly clear health implications, the human microbiome has become an obvious target of amplicon sequencing experiments (Gupta et al., 2019; Jovel et al., 2016). We know much less about microbial communities in other organisms, often differing substantially from human microbiomes in their taxonomic composition, diversity patterns, and biological roles. Insect microbiomes are one such category. In the era of the dramatic decline of insect biodiversity, rightfully called the insect apocalypse (Sánchez-Bayo & Wyckhuys, 2019; Wagner et al., 2021) within the ongoing sixth mass extinction (Cowie et al., 2022; Wagner et al., 2021), it is crucial to describe and understand the potential role of symbionts in insects’ adaptation to changing environments. We know that insect-associated bacteria are taxonomically and functionally diverse and highly significant from the evolutionary and ecological perspective, and some of them are extremely divergent from the closest relatives known from other environments. Unfortunately, outside a few model species and a few symbiont clades, our knowledge of the diversity and distribution of insect microbial communities is limited. Although recent decades have seen a boom in molecular techniques, we still do not have labour- and cost-effective tools to comprehensively describe symbionts’ spatio-temporal dynamics at the level of whole insect communities.

Here, we presented our proof of concept, showing that amplicon sequencing is a powerful approach for surveying the composition of the microbial community and the bacterial absolute abundance of non-model insects. The proposed protocol is not only cost-effective and time-efficient but also highly versatile. Our custom DNA extraction method, costing just $0.75 per sample, is significantly cheaper and more efficient than commercial kits, enabling the processing of three times as many samples within the same time frame. Given the demonstrated lack of a significant difference in amplification efficiency, we strongly recommend incorporating this extraction protocol into laboratory pipelines targetting insect microbiome analyses. Our results clearly demonstrate that using the proposed protocol, we have successfully amplified different regions of bacterial 16S rRNA alongside the insect COI marker gene. The versatility of our approach makes it relatively easy to implement additional marker genes for specific research questions. For instance, we can successfully implement amplicon-based multitarget genotyping involving fungal or protist taxa, enabling a comprehensive analysis of microbiota diversity. Moreover, incorporating plant-specific genes into the workflow could facilitate the reconstruction of insect diets or pollination networks. Perhaps most importantly, our pipeline enables reliably reconstructing bacterial absolute abundances as a “byproduct” of bacterial metabarcoding. With the significant decrease in the cost of high-throughput sequencing observed in recent years (Hu et al., 2021), the number of publications tackling insect microbiomes has increased. However, the vast majority of such publications use relative abundances, and outside of a few notable exceptions (Hammer et al., 2017; Ravenscraft et al., 2019; Sanders et al., 2017a; Surmacz et al., 2024), we virtually know nothing about the microbiome absolute abundances in insects (Hammer et al., 2019; Sanders et al., 2017b). Noteworthy, our amplicon-based quantification is congruent with results described by other scientists using well-established tools such as qPCR for *A. mellifera* (Hammer et al., 2019). Here, for the first time to our knowledge, we describe not only microbiome composition for the whole natural insect community but also provide information about the variation of bacterial quantities within and between species. The possibility of quantifying microbial abundances alongside their composition opens up entirely new avenues for understanding insect-symbiont interactions. Although implementing a multitarget amplicon-based survey has significant advantages that have already been addressed, it is not free of serious caveats. Those caveats need to be carefully considered and addressed if the method is to provide reliable insights into host-microbe associations. Firstly, the capability of relatively short gene fragments to reflect an actual microbiome composition. We showed that each of the V3-V4 and V4 regions of the 16S rRNA gene were able to detect the majority of specialized insect symbionts present in the metagenome. However, in line with the previous research on various model organisms and environmental systems (Bobay et al., 2020; De Filippis et al., 2017), our data suggests that results obtained from amplicon sequencing need to be carefully considered and interpreted, as depending on targeted region and specific primer pair, they may not necessarily reflect an actual microbial pattern. Secondly, authors describing insect microbiomes are usually unaware of the numerous sources of contamination throughout the laboratory process. Although the necessity of using various negative controls has been brought up in high-impact publications (e.g. Knight et al., 2018), most authors do not mention the use of negative controls or limit themselves to describing the lack of a band on electrophoresis gels. Our results clearly show that even without a visible band on a gel, the proportion of PCR or extraction contaminants can blur the image of microbiome composition, especially in species with virtually no microbiome (Fig. 6)

**Fig 6.**
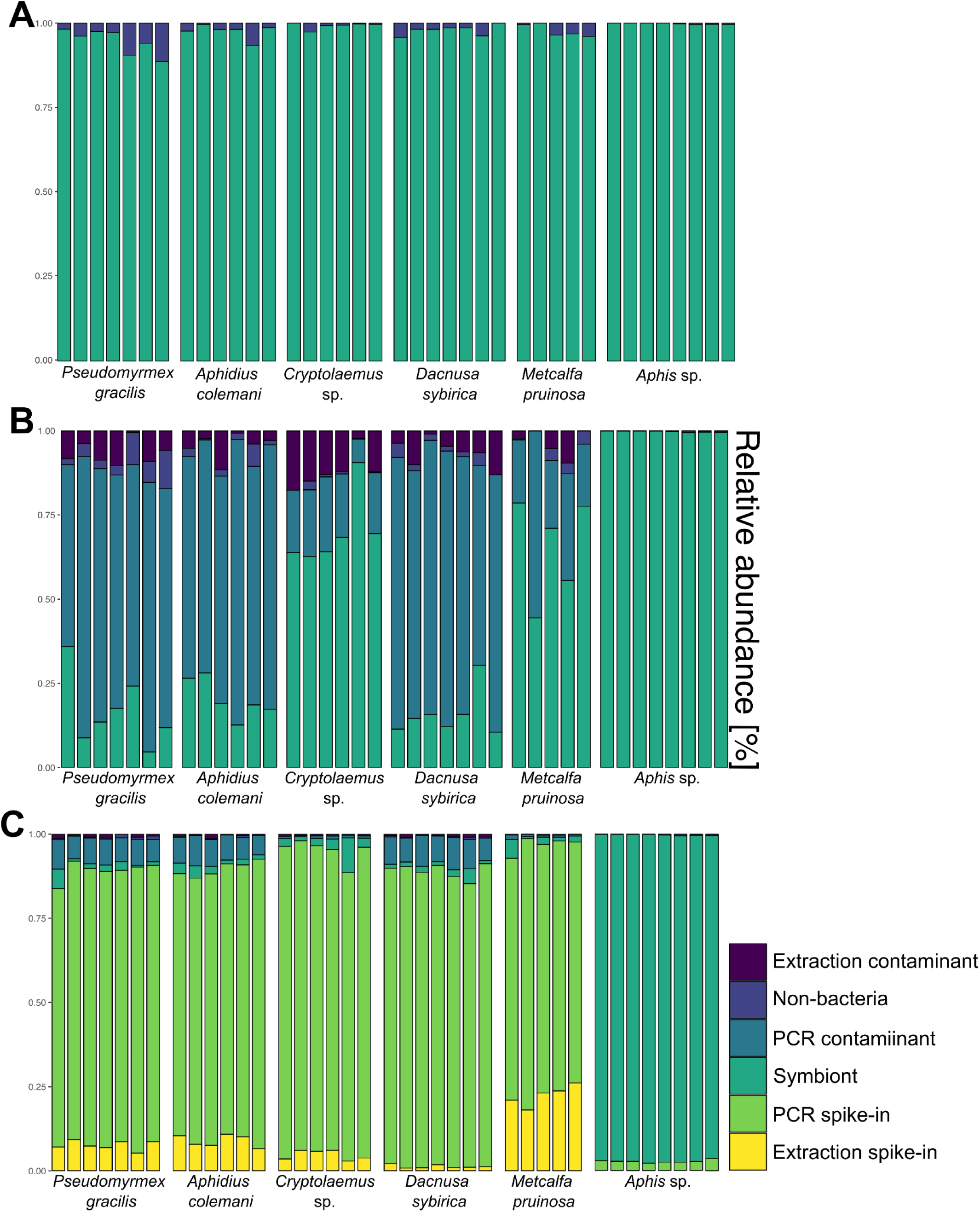
The visualization of the decontamination and quantification of example bacterial 16S rRNA sample data. A. Usually, most pipelines finish at a stage of taxonomy assignment with recognition of non-bacterial reads (Archea, Mitochondria, Chloroplasts, etc.), and after deleting them, scientists proceed with the various analyses. B. We can recognise contaminants of different origins by introducing decontamination based on numerous negative controls (extraction and PCR). C. When we implement quantification based on the count ratio of artificial constructs added during DNA extraction and PCR steps to the real bacterial reads, we obtain absolute quantity estimates for the 16S targets present in different insects.

Interestingly, one of the bacterial OTUs present in all data generated in our Lab, assigned by our decontamination script as PCR contaminant and taxonomically classified as *Brachybacterium;* has been described by numerous authors as a microbiome member of: coleopterans (Ali et al., 2019), rice phyllosphere (Wiraswati et al., 2023), or defensive skin symbiont of amphibians (Bletz et al., 2017). Comparing the sequence of our contaminant with sequences from the mentioned studies, we obtained 100% support for query coverage and sequence identity. Naturally, we cannot exclude the possibility that this particular bacteria is widespread and can play an important function across different organisms. However, the lack of sequenced negative controls in the abovementioned examples can indicate this OTU as a characteristic contaminant of PCR Master Mix. The importance of negative controls has been demonstrated in the recent publication from our Group (Surmacz et al., 2024). By reanalyzing published data about the Tardigrade microbiome (Mioduchowska et al., 2023) and generating a new dataset with a strict usage of various negative controls, our colleagues showed how improper experiment design can lead to erroneous conclusions about the composition and potential role of the microbiome.

We demonstrated the utility and versatility of our approach on the whole insect community from NE Greenland, capturing ca. 50% of its species and reconstructing their microbiome compositions and quantities. The more traditional techniques for untangling relationships between hosts and their symbionts on a bigger scale presented significant obstacles. Traditional species delimitation by experts in the field is time-consuming and requires a vast network of connections. Similarly, extracting DNA by commercial kits and generating marker gene sequences separately for insects and their microbes is both time- and cost-insufficient. To put that into perspective, laboratory processing to generate COI barcodes for coleopterans and *Wolbachia* MLST-typing for approximately 300 species (ca. 1200 specimens) using Sanger sequencing to reconstruct their relationships took around three years (Kajtoch et al., 2019). We addressed similar hypotheses using our approach by processing 1842 Phorid fly specimens in less than a year (Nowak et al., 2024). Hence, the presented pipeline has the potential to revolutionize the field of insect-microbiome studies, providing a needed tool for addressing bold ecological questions. We can track insects and their microbiome dynamics almost in real-time, without breaking the bank, which is extremely important during an ongoing biodiversity crisis.

## 5. Data availability

All the generated sequencing data will be available in the GenBank upon publication.

Due to the size of the file Supplementary Table 13 is available under the link https://drive.google.com/file/d/1uzKYeTeqp0JyMWEkUT9LxFaKrdVrp2Mo/view?usp=drive_link

## 6. Code availability

Bioinformatic pipelines are available on the GitHub repository: https://github.com/Symbiosis-JU/Proof_of_Concept

## 7 Acknowledgements

The project was supported by the Polish National Agency for Academic Exchange grant PPN/PPO/2018/1/00015 (to P.Ł.), Polish National Science Centre grants: 2018/31/B/NZ8/01158 (to P.Ł.), 2018/30/E/NZ8/00880 (to P.Ł.), 2017/26/D/NZ8/00799 (to A.M.) and has received funding from INTERACT III Transnational Access under the European Union H2020 Grant Agreement No. 629.

---

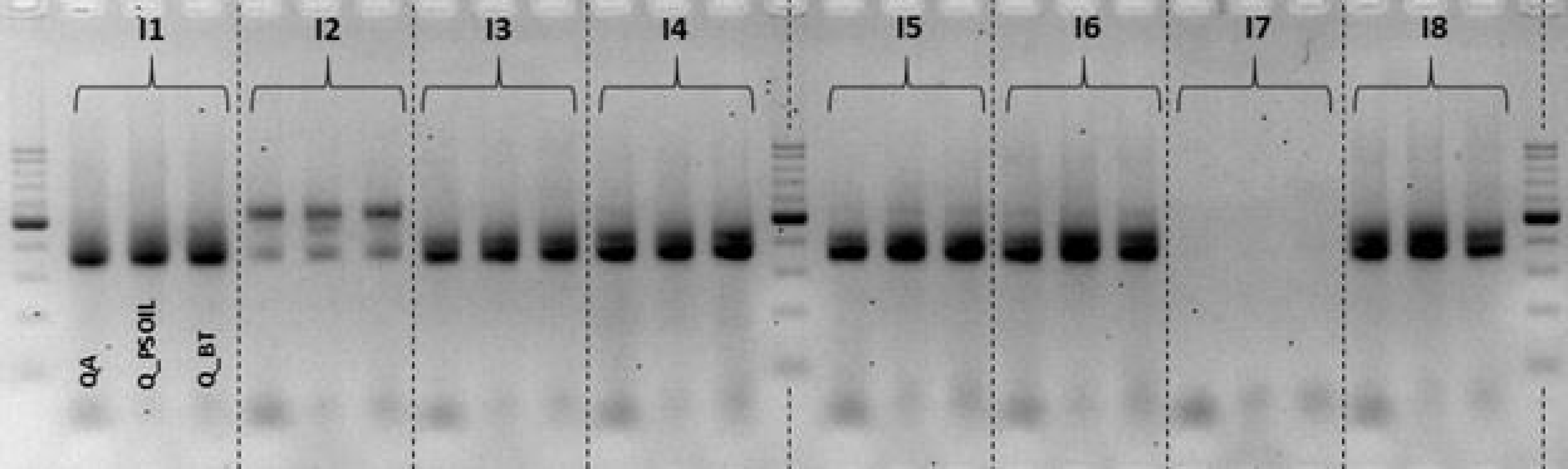

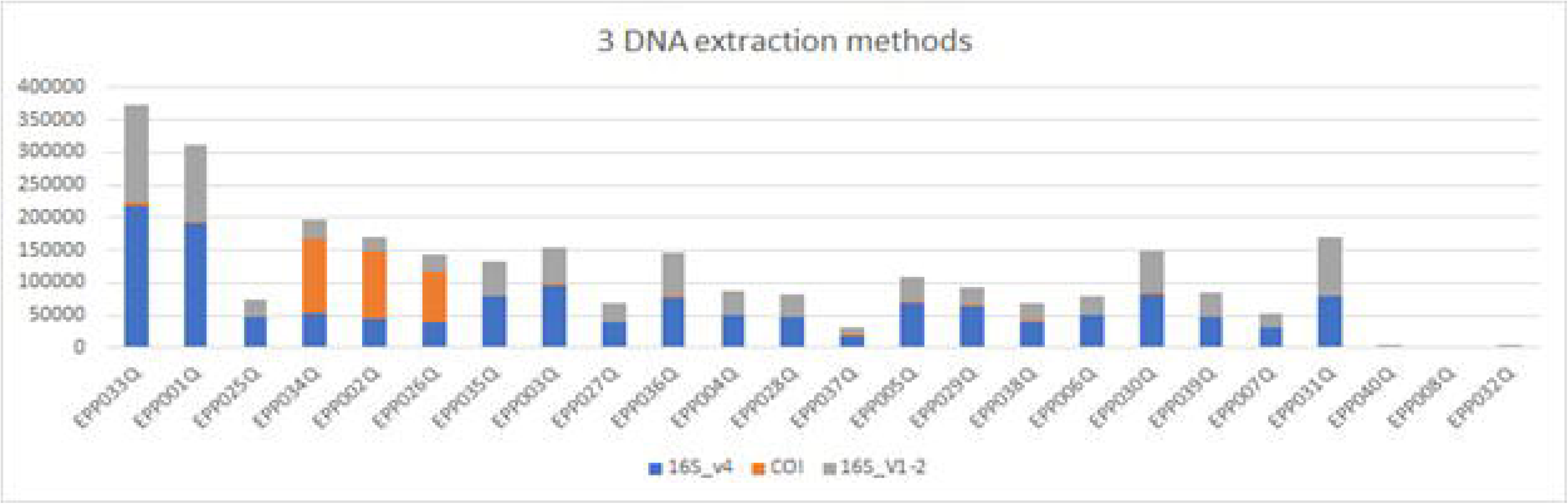

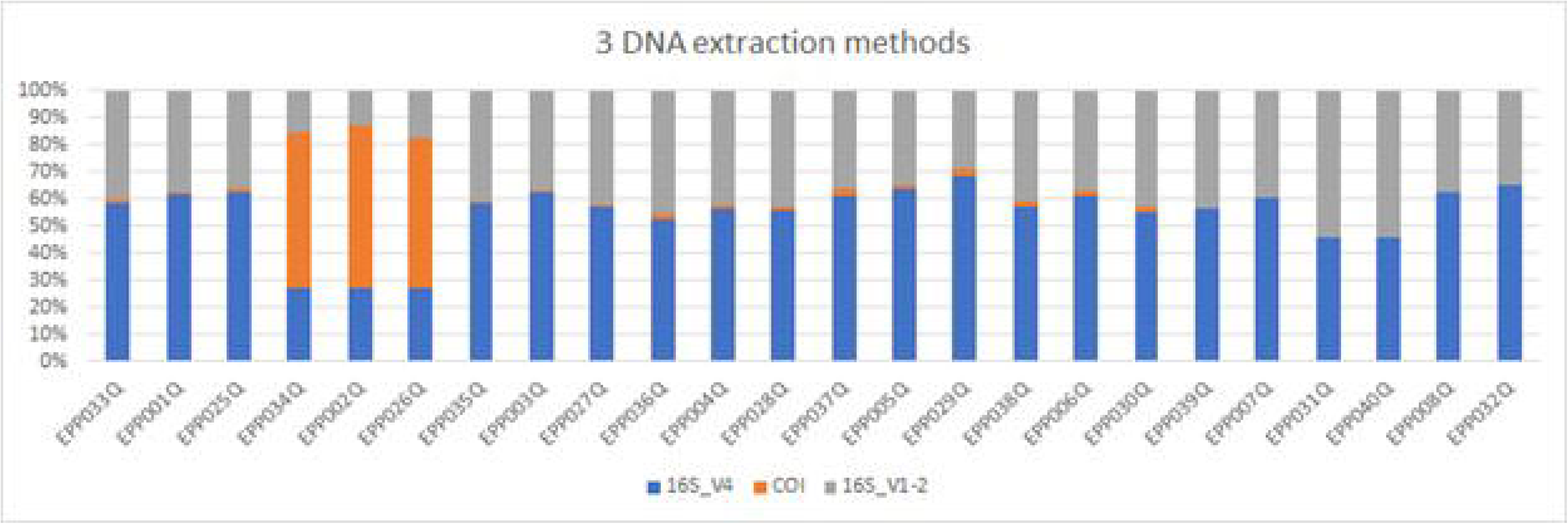

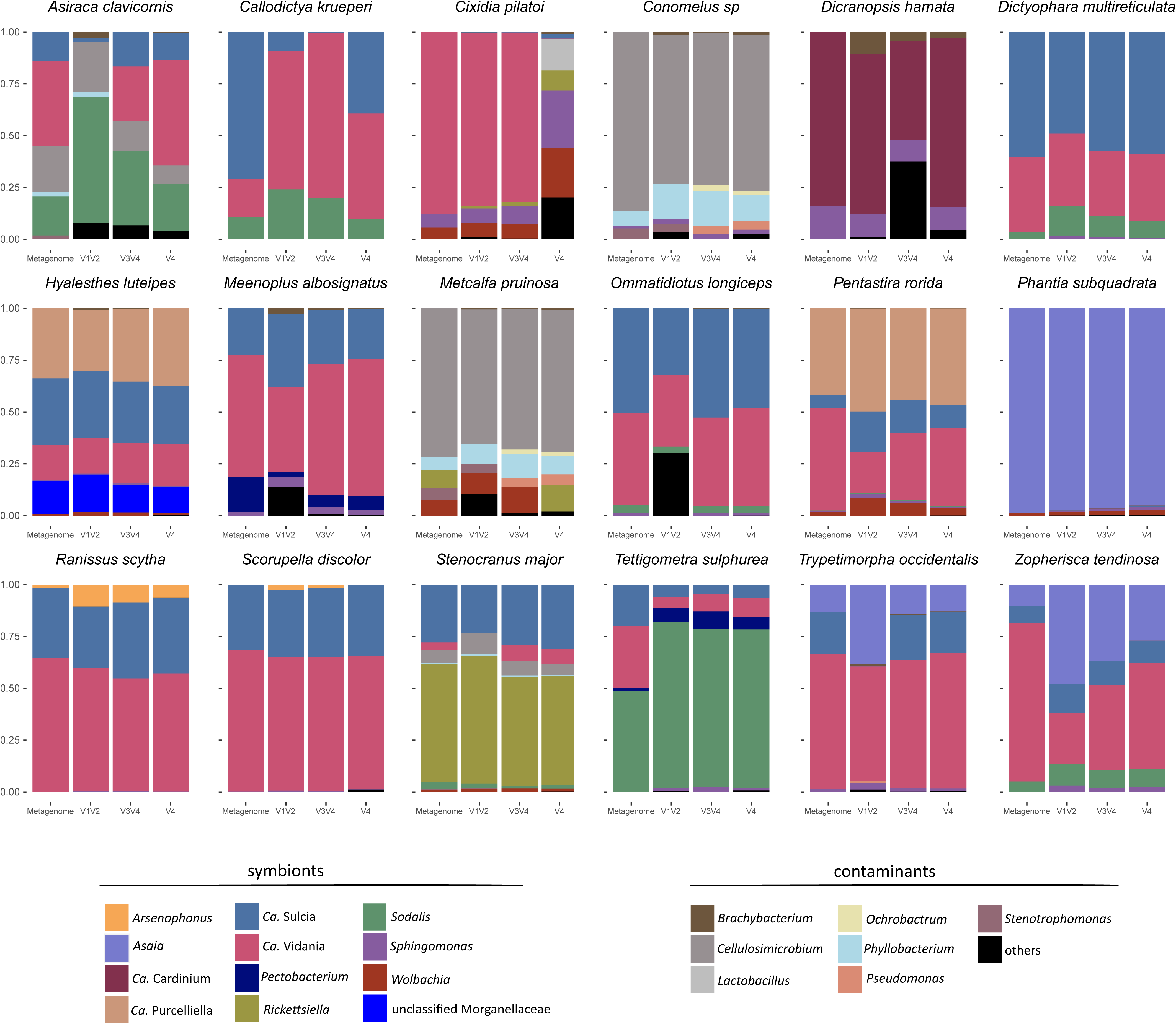

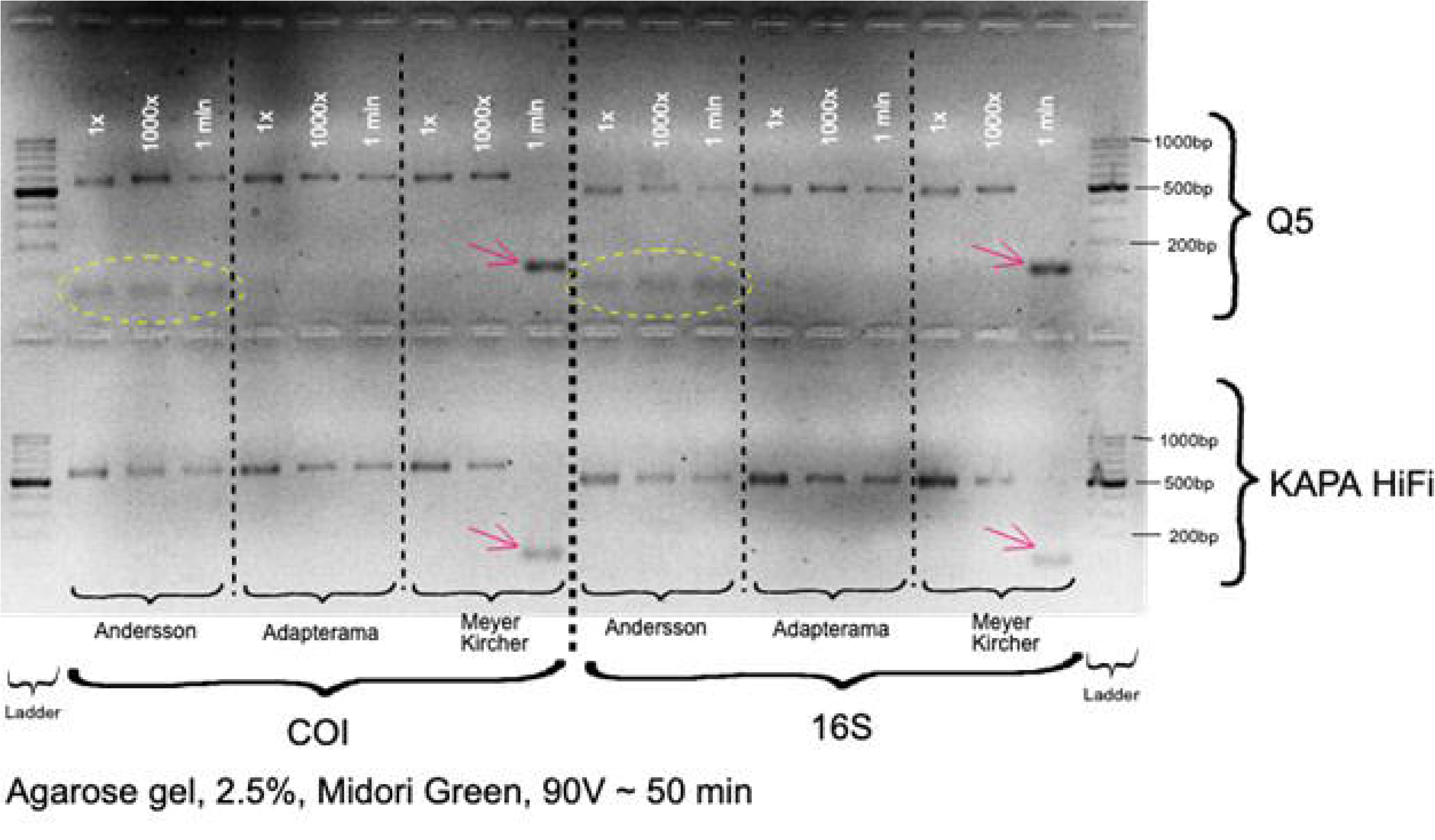

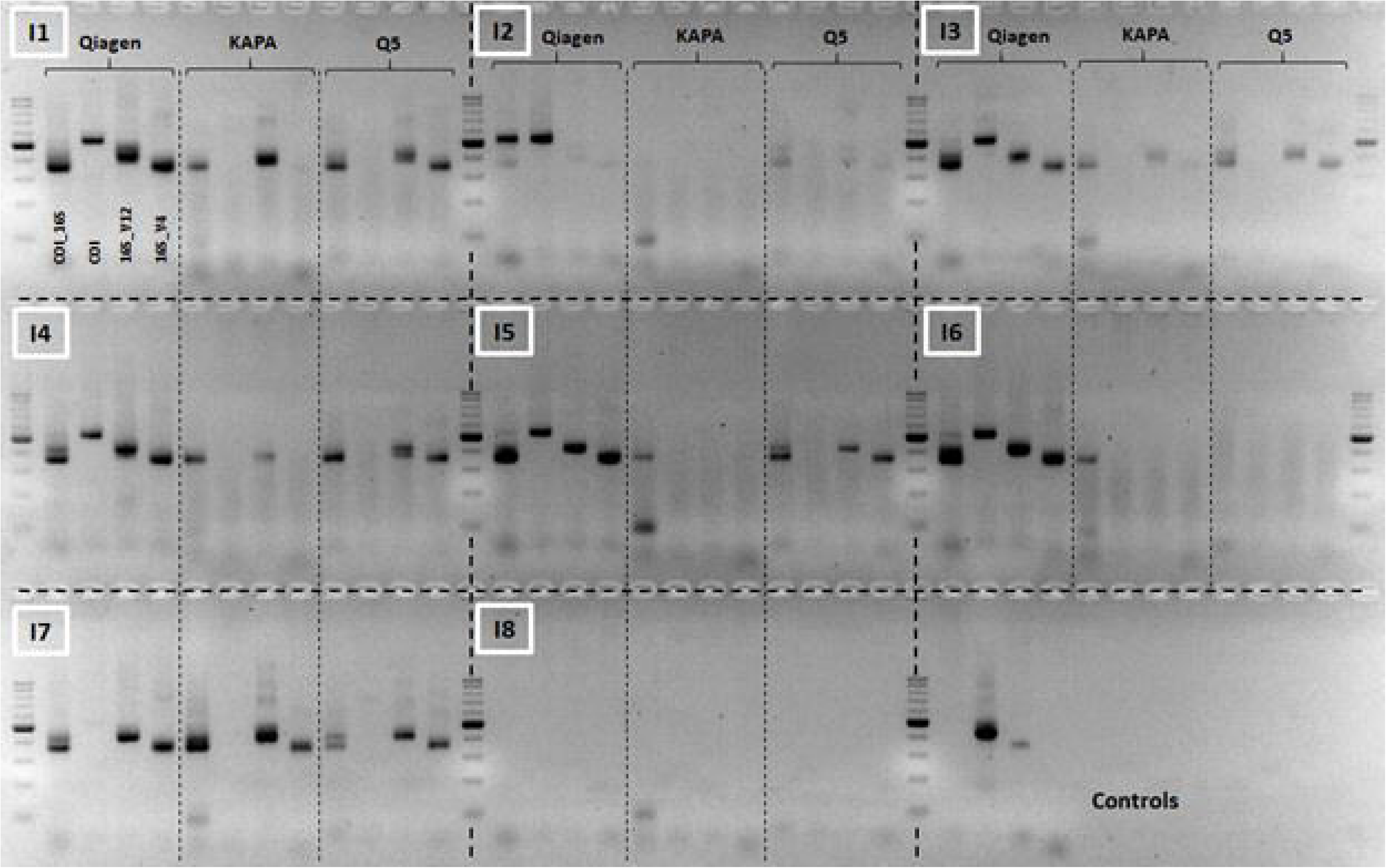

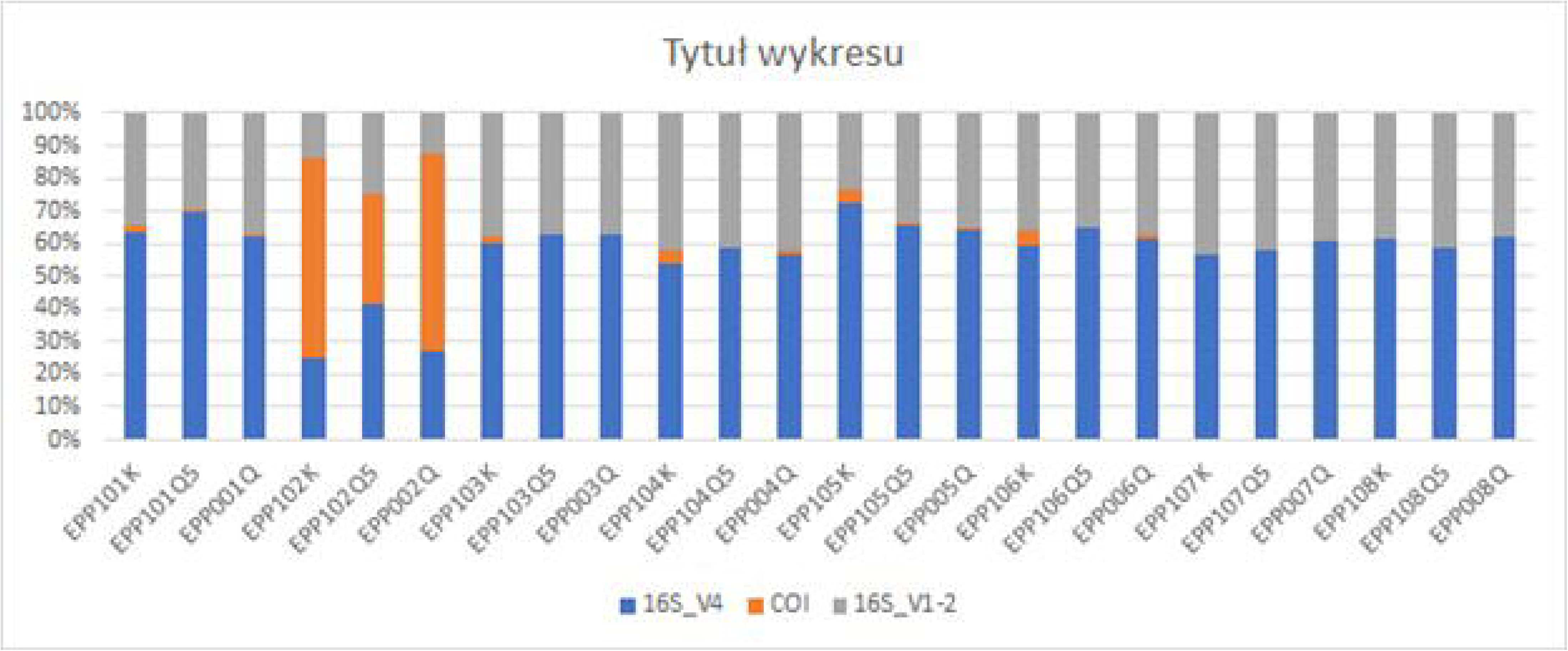

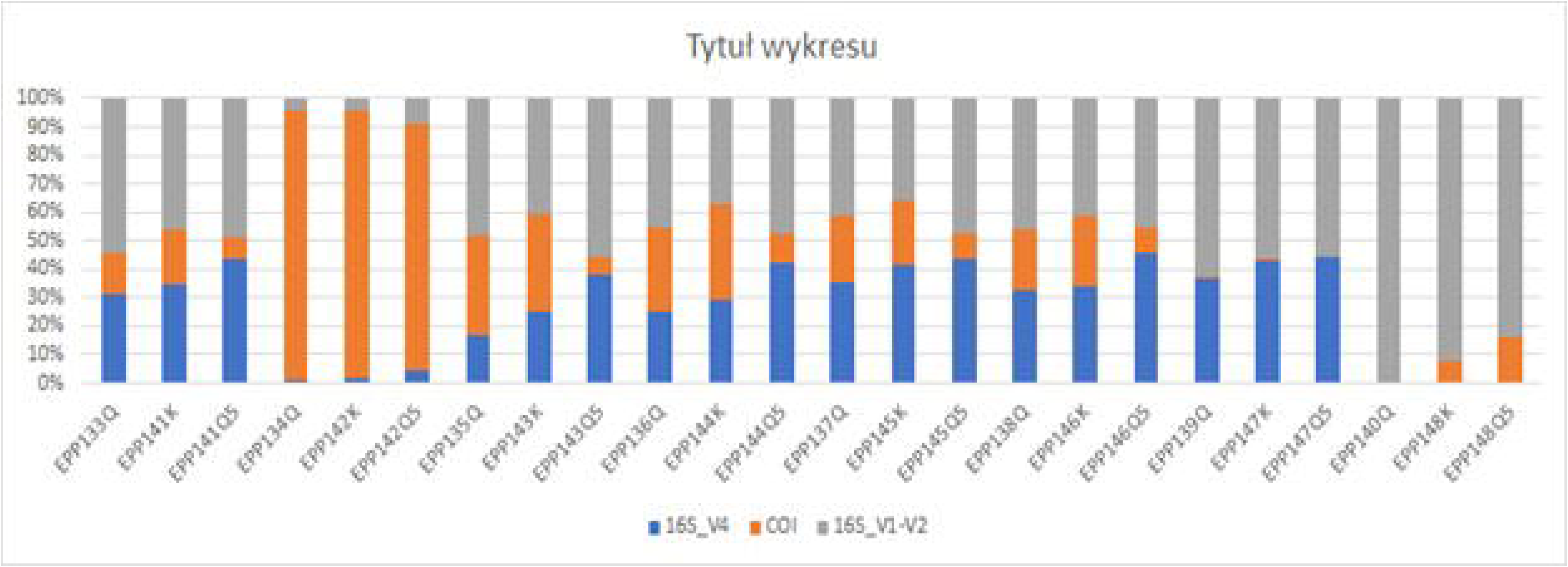

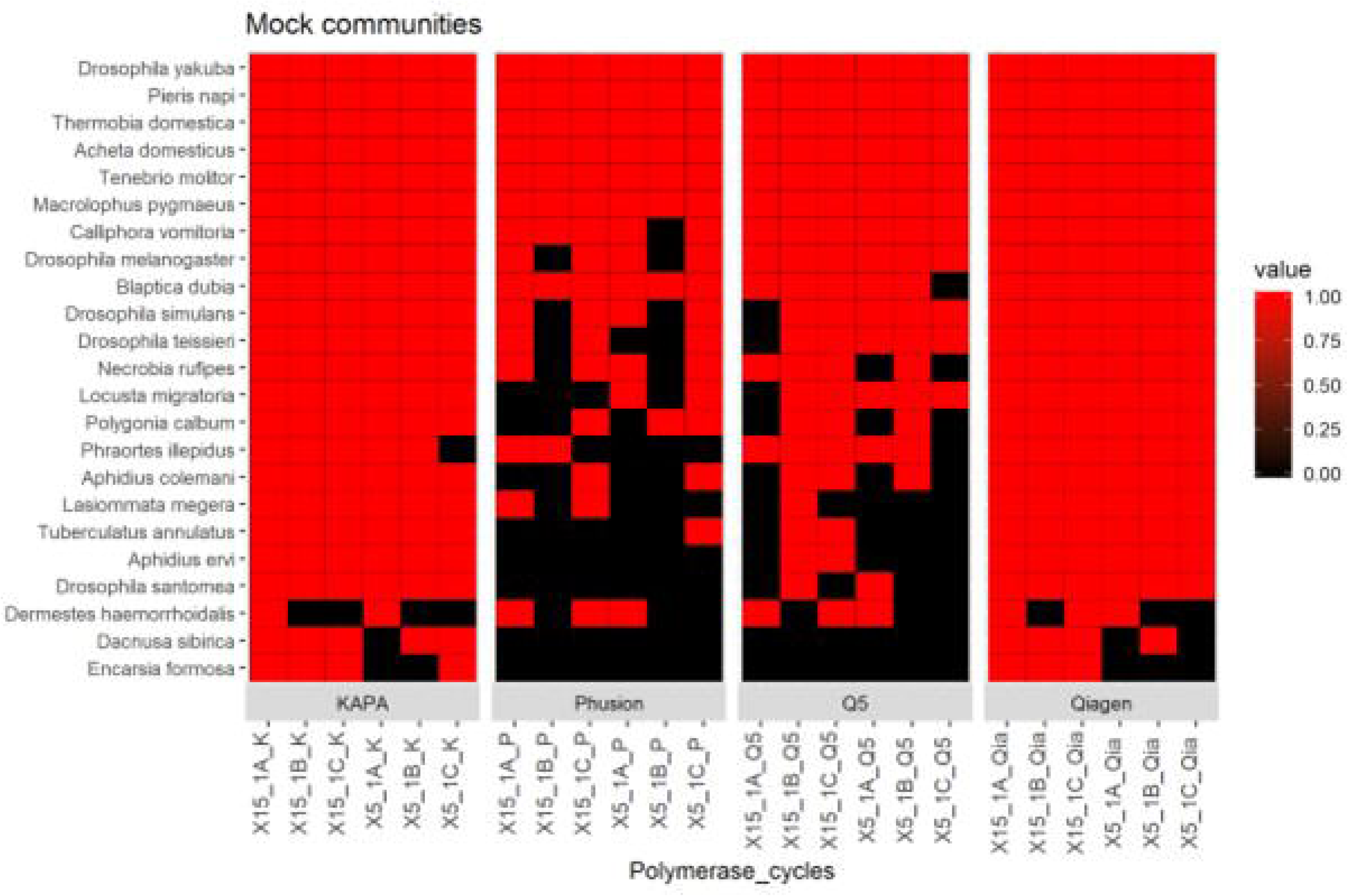

